# KSR1 mediates small-cell lung carcinoma tumor initiation and cisplatin resistance

**DOI:** 10.1101/2024.02.23.581815

**Authors:** Deepan Chatterjee, Robert A. Svoboda, Dianna H. Huisman, Benjamin J Drapkin, Heidi M. Vieira, Chaitra Rao, James W. Askew, Kurt W. Fisher, Robert E. Lewis

## Abstract

Small-cell lung cancer (SCLC) has a dismal five-year survival rate of less than 7%, with limited advances in first line treatment over the past four decades. Tumor-initiating cells (TICs) contribute to resistance and relapse, a major impediment to SCLC treatment. Here, we identify Kinase Suppressor of Ras 1 (KSR1), a molecular scaffold for the Raf/MEK/ERK signaling cascade, as a critical regulator of SCLC TIC formation and tumor initiation *in vivo*. We further show that KSR1 mediates cisplatin resistance in SCLC. While 50-70% of control cells show resistance after 6-week exposure to cisplatin, CRISPR/Cas9-mediated KSR1 knockout prevents resistance in >90% of SCLC cells in ASCL1, NeuroD1, and POU2F3 subtypes. KSR1 KO significantly enhances the ability of cisplatin to decrease SCLC TICs via *in vitro* extreme limiting dilution analysis (ELDA), indicating that KSR1 disruption enhances the cisplatin toxicity of cells responsible for therapeutic resistance and tumor initiation. The ability of KSR1 disruption to prevent cisplatin resistant in H82 tumor xenograft formation supports this conclusion. Previous studies indicate that ERK activation inhibits SCLC tumor growth and development. We observe a minimal effect of pharmacological ERK inhibition on cisplatin resistance and no impact on TIC formation via *in vitro* ELDA. However, mutational analysis of the KSR1 DEF domain, which mediates interaction with ERK, suggests that ERK interaction with KSR1 is essential for KSR1-driven cisplatin resistance. These findings reveal KSR1 as a key regulatory protein in SCLC biology and a potential therapeutic target across multiple SCLC subtypes.

**Statement of Implication:** Genetic manipulation of the molecular scaffold KSR1 in small-cell lung cancer cells reveals its contribution to cisplatin resistance and tumor initiation.

## Introduction

Small-cell lung carcinoma (SCLC) is an aggressive, lethal, and highly metastatic disease accounting for 13% of all lung cancers^(1)^. Predominately affecting heavy smokers, SCLC is also highly metastatic with 50%–80% of patients harboring metastases at the time of autopsy^(2)^. SCLC was categorized as a recalcitrant cancer in 2012 due to only modest improvements in SCLC detection, therapy, and survival over the past 40 years^(1)^. The five-year relative survival rates for patients with local, regional, and distant SCLC are 27%, 16%, and 3%, respectively (Cancer.net, ASCO, 2022). Currently, SCLC tumors are treated with first line therapy (cisplatin or carboplatin plus etoposide chemotherapy, anti-PDL1 antibody, atezolizumab), second line (topotecan, lurbinectedin^(3)^), and for some patients thoracic radiation therapy^(4)^. Although SCLC tumors are responsive to therapy initially, residual disease quickly develops resistance leading to the low five-year survival^(5)^. Despite extensive genomic and molecular analysis, few biochemical pathways in SCLC have yielded convincing therapeutic targets that might selectively disable SCLC with minimal toxicity to normal tissue. Identification of functionally significant pathways and proteins regulating growth and survival whose inhibition selectively disables the tumor’s self-renewal capacity, could lead to the development of novel therapeutic strategies for SCLC, especially promoting a more durable treatment response and lesser incidences of recurrence.

Rigorous and innovative basic science using state-of-the-art genetically engineered mouse models (GEMM), and extensive cell lines painstakingly generated by multiple labs have led to key discoveries regarding the cells-of-origin and the common recurring mutations that underlie SCLC^(6–13)^. These discoveries have yielded comprehensive genomic profiles and a durable classification of SCLC subtypes based on the differential expression of four key transcription factors, ASCL1 (SCLC-A), NeuroD1 (SCLC-N), POU2F3 (SCLC-P) and Yap1 (SCLC-Y)^(10,14)^, which has been reclassified recently as undifferentiated SMARCA4-deficient malignancies^(15)^. Based on PDX studies, a fifth subtype based on the bHLH proneural transcription factor ATOH1, has also now been proposed^(16,17)^. A subpopulation of self-renewing pulmonary neuroendocrine cells (PNECs) can give rise to SCLC-A tumors when TP53, Rb, and Notch mutations cause constitutive activation of stem cell renewal and altered deprogramming^(18)^. The transformed, self-renewing population within SCLC-A, with a neuroendocrine origin, is characterized by high levels of CD24 and EPCAM and low levels of CD44^(19)^. Myc overexpression, which is common in aggressive SCLC, drives conversion of SCLC-A to SCLC-N^(20)^. Expression of NeuroD1 in SCLC-N is linked to the development of metastases and aggressive SCLC^(21)^. MYC alternatively promotes POU2F3^+^ tumors from a distinct cell type, potentially of a chemosensory lineage known as Tuft cells^(22–24)^.

SCLC tumors have a small population of tumor initiating cells (TICs) essential for the initiation, long-term propagation, and metastatic ability of the tumor^(19)^. In contrast, SCLC tumors also have a bulk non-TIC population that is highly proliferative but inefficient at establishing new tumors *in vivo*^(19)^. TICs may be the sanctuary population within the bulk tumor responsible for therapeutic resistance in SCLCs^(25–31)^. Recent studies have revealed that a subpopulation of drug-tolerant cells known as ‘persisters’ plays a pivotal role in the development of drug resistance within tumors. Like TICs, persisters often exhibit the ability to maintain their viability during therapy, albeit at a slower rate or in a dormant state. This observation aligns with the accumulating evidence suggesting a shared identity between TICs and DTPs^(32)^. Despite the negative implication of TICs/DTPs in patient survival^(33,34)^, very little is understood about the underlying mechanisms that support their formation and function.

Kinase suppressor of Ras (KSR) 1 and KSR2 are molecular scaffolds that promote extracellular signal-regulated kinase (ERK) signaling through the mitogen-activated protein kinase (MAPK) pathway but have differing expression profiles and physiological functions^(35,36)^. KSR1 is required for oncogenic Ras-induced transformation and enhances the extent and duration of ERK activation by growth factor receptor tyrosine kinases, but it is dispensable for normal cell survival^(37–39)^. *Ksr1^−/−^* knockout mice are fertile, and developmentally normal, but are resistant to Ras-driven tumor formation^(40)^. KSR1 regulates Ras-dependent transformation and survival through both the Raf/MEK/ERK kinase cascade^(37,38,41–44)^ and AMPK ^(35,41,45)^. KSR1 depletion suppresses transformation in Ras mutated colorectal cancer (CRC) cell lines and prevents drug-induced TIC upregulation in Ras mutant colon and non-small lung cancer cell lines ^(46,47)^. These observations suggest that KSR1 and the downstream effectors it modulates are essential for tumor formation but dispensable for normal tissues. Targeting these effectors should be selectively toxic to tumors. The closely related member of KSR1, Kinase Suppressor of Ras 2 (KSR2) is an essential regulator of clonogenicity and self-renewal in SCLC-A (submitted) (bioRxiv 2022.02.11.480157v4), which led us to study the function of KSR1 in SCLC, across all the subtypes. Our work reveals KSR1 regulates clonogenicity and tumor initiation in SCLC, both *in vitro* and *in vivo*. It also highlights KSR1 as a novel player in promoting therapy resistance in these cells. This result defines a novel mechanism of tumor initiation in SCLC and a potential therapeutic vulnerability.

## Materials and Methods

### Cell culture

Human small-cell lung carcinoma (SCLC) cell lines NCI-H82 (H82), NCI-H524 (H524), HCC33, NCI-H209 (H209), and NCI-H2107 (H2107) were gifts of John Minna (UT Southwestern). HEK293T, other human small-cell lung carcinoma cell lines NCI-H526 (H526) and NCI-H1048 (H1048) and the human non-small cell lung cancer (NSCLC) cell line, A549 were purchased directly from ATCC. The SCLC cell lines H82, H524, HCC33, H209, H526 and the NSCLC cell line A549 were cultured in RPMI 1640 (VWR #16750-106) medium containing 10% fetal bovine serum (FBS). The SCLC cell lines H2107 and H1048 were cultured in HITES medium (ATCC #30-2006) containing 5% fetal bovine serum (FBS). All the cells were grown at 37°C with ambient O_2_ and 5% CO_2_. No further authentication of cell lines was performed by the authors. Cells were routinely tested for mycoplasma using the 16S rRNA gene detection method by isothermal PCR (MycoStrip® Mycoplasma detection kit Invivogen #rep-mys-50) and confirmed to be free of contamination, with the most recent test conducted on 12.18.2024.

### Generation of pooled genomic KSR1 knockout (KO) cell lines H82, H209, H2107, H526, H1048

Single guide RNA (sgRNA) sequences targeting KSR1 or non-targeting control (NTC) were inserted into pLentiCRISPRv2GFP (Addgene #82416). The constructs were polyethylenimine (PEI) (Tocris #7854) transfected into HEK293T cells along with psPAX2 lentiviral packaging construct (Addgene #12260) and pMD2.G envelope construct (Addgene #12259). Lentivirus-containing media was harvested at 72-hour post-transfection and used to infect the H82, H209, H2107, H526 and H1048 cells with 8 µg/mL Polybrene (Sigma #TR1003G). GFP+ cells were selected by fluorescence-activated cell sorting after 72-96 hours on a BD FACSAria II using FACSDiva software.

### Generation of inducible genomic KSR1 KO cell lines

sgRNA targeting KSR1 or non-targeting control (NTC) sgRNA were inserted into TLCV2-RB1 (pLentiCRISPRv2GFP modified with a puromycin selection marker) (Addgene #87836). The constructs were PEI transfected into HEK293T cells along with psPAX2 lentiviral packaging construct (Addgene #12259) and pMD2.G envelope construct (Addgene #12259). Lentivirus containing media was harvested at 72-h, and added to H82, H524, and HCC33 cells with polybrene. 48-hour post infection, cells were selected in 2 µg/mL Puromycin (Invivogen #antpr1). The cells were then treated with 2 µg/mL doxycycline (Sigma #D9891) for 72 hours, inducing Cas9-2A-eGFP. GFP+ cells were finally selected by fluorescence-activated cell sorting.

### Generation of ectopically expressed KSR1 construct in inducible genomic KSR1 KO cell line H82, and wild-type H209 and H2107

A murine KSR1 cDNA^(47)^ resistant to recombination by sgRNA sequences targeting human KSR1 was cloned into MSCV-IRES-KSR1-RFP (Addgene #33337) and PEI transfected into HEK293T cells along with pUMVC retroviral packaging construct (Addgene #8449) and CMV5-VSVG envelope construct (Addgene #8454). Retrovirus containing media was harvested at 96-hour and added to inducible KSR1 KO H82, and wild-type H209 and H2107 cells with 8 mg/ml polybrene. RFP-selection was performed by fluorescence-activated cell sorting 48 h post-infection.

### Generation of KSR1 KO H82 cell line expressing ERK-binding deficient KSR1

A KSR1 transgene deficient in binding ERK due to engineered mutation in the DEF-domain, KSR1^AAAP^ ^(37,42)^ was cloned into MSCV-IRES-KSR1-RFP and PEI transfected into HEK293T cells along with pUMVC retroviral packaging construct (Addgene #8449) and CMV5-VSVG envelope construct (Addgene #8454). Retrovirus-containing media was harvested at 96 hours and added to KSR1 KO cells with 8 mg/ml polybrene. RFP-selection was performed by fluorescence activated cell sorting 48 hours post-infection.

### Cell lysis and Western blot analysis

Whole cell lysate was extracted in radioimmunoprecipitation assay (RIPA) buffer containing 50 mM Tris-HCl, 1% NP-40, 0.5 % Na deoxycholate, 0.1 % Na dodecyl sulfate, 150 mM NaCl, 2 mM EDTA, 2 mM EGTA, and protease and phosphatase inhibitors aprotinin (0.5 U/ml), leupeptin (20 mM), and NA_3_VO_4_ (0.5 mM). Protein concentration was estimated using BCA protein assay (Promega #PI-23222, PI-23224). Samples were diluted using 1X sample buffer (4X stock, LI-COR Biosciences #928–40004) with 100 mM dithiothreitol (DTT) (Sigma #D9779-5G). Protein was separated using 8–12% SDS-PAGE and transferred to nitrocellulose membranes. The membrane was blocked with Odyssey TBS blocking buffer (LICOR Biosciences #927–50003) for 45 min at room temperature, then incubated with primary antibodies at least overnight at 4°C. IRDye 800CW and 680RD secondary antibodies (LI-COR Biosciences # 926–32211, # 926–68072) were diluted 1:10,000 in 0.1% TBS-Tween and imaged on the Odyssey Classic Scanner (LI-COR Biosciences).

### Colony Formation Assay

For colony formation assays, SCLC cells were dissociated by gentle pipetting. Cells were DAPI (Abcam #ab228549) stained for viability determination and live GFP+/RFP+ cells were single cell sorted as one cell per well into a 96-well plate (Fisher #12556008) filled with complete media (200 μl/well). 50 µl fresh media was added to the wells once every week. Three to four weeks later, clonogenicity was assessed by measuring luminescence using CellTiter-Glo 2.0 reagent (Promega #G9242) and POLARstar Optima plate reader according to the manufacturer’s protocol. CellTiter-Glo® readings greater than 300 Relative Luminescence Units (RLUs) in colony forming assays were considered colonies.

### Extreme Limiting Dilution Analysis (ELDA)

#### In vivo

The viable cell number was assessed by replicate cell counts on a hemocytometer using Trypan Blue exclusion. Viable cell number was used to derive a titration of cell numbers for implantation. Cells were diluted in 50 μl media (RPMI +10% FBS) and 50 μl Cultrex PathClear BME (Trevigen # 3632-005-02). Cells were injected subcutaneously into the shoulders and flanks of six eight-week-old *Prkdc^em26Cd52^Il2rg^em26Cd^*^22^/NjuCrl (NCG) mice. Three replicates for each dilution were used. Injection sites were palpated biweekly to monitor for tumor growth and all mice sacrificed when any one tumor reached 1 cm. Tumors that formed were scored as 1 and the absence of tumor formation was scored as 0 for the extreme limiting dilution analysis (ELDA). Tumors that formed were analyzed for absence/presence of KSR1 by Western blot.

#### In vitro

Cells were seeded in 96-well plates (Fisher #12556008) at decreasing cell concentrations (1000 cells/well – 1 cell/well) at half log intervals (1000, 300, 100, 30, 10, 3, 1), 12 wells per condition except for the 10 cells/well condition, for which 24 wells were seeded. Cells were cultured for 10-15 days, and wells with spheroids >100 µM were scored as spheroid positive. To assess the effect of cisplatin on TIC frequency, cells were plated in the presence of the indicated dose of cisplatin (Sigma #232120) and scored as positive/negative after 10-15 days as previously described. To assess the effect of ERK inhibition on TIC frequency, cells were plated in the presence of 2 µM SCH772984 (SelleckChem #S7101) and scored as positive/negative after 10-15 days as previously described. TIC frequency from *in vitro* and *in vivo* analyses and the significance between groups was calculated using ELDA software https://bioinf.wehi.edu.au/software/elda/^(48)^.

### Dose-Response Curve

Cells were seeded at 10,000 cells per well in 50 µl in the inner-60 wells of 96-well black-sided, clear-bottom culture plates (Fisher #237105) and allowed to incubate for 24 hours prior to drug treatment. After 24 hours, cells were treated with nine different doses of drug concentrations spanning a range of concentrations (20, 100, 200, 500, 1000, 2000, 10000, 20000, 50000 nM) with each column of the 96 well plate getting a particular concentration of drug for 72 hours prior to assessment of cell viability using CellTiter-Glo® 2.0 reagent. Data were analyzed by non-linear regression using GraphPad Software, Inc.

### Growth Curve

Cells were seeded at low density (500 cells/well) in 100 µl in replicate 96-well plates (Fisher #237105), with each condition plated in a single column of the 96 well plate. Each replicate plate was reserved for a single day. Wells were fed with complete media and assessed daily for growth for a period of 9-15 days, through assessment of cell viability using CellTiter-Glo® 2.0 reagent. Data were analyzed by non-linear regression using GraphPad Software, Inc.

### *In vivo* Tumor Volume Curve

The viable cell number was assessed by replicate cell counts on a hemocytometer using Trypan Blue exclusion. Viable cell number was used to derive a dilution of 2.5 million cells for implantation. Cells were diluted in 50 μl media (RPMI +10% FBS) and 50 μl Cultrex PathClear BME (Trevigen # 3632-005-02). NTC or KSR1 KO H82 cells were injected subcutaneously into the left flank of twenty-four eight-week-old *NUJ (nude)* mice. Twelve replicates for each condition were used. Injection sites were palpated daily to monitor for tumor growth. Six days after injection of the cells, mice with NTC or KSR1 KO tumors were evenly divided into vehicle (PBS) treatment and cisplatin (5 mg/kg by intraperitoneal (i.p.) injection, twice a week) treatment. Cisplatin was a gift from Michael Kareta (Sanford Research). Primary tumor growth was measured and quantified using calipers. Mice were sacrificed, once tumors reached 1000 mm^3^. Data were analyzed by non-linear regression using GraphPad Software, Inc. All procedures conformed to accepted practices for the humane use of experimental animals and were approved by the UNMC Institutional Animal Care and Use Committee.

### Cell Cycle Analysis

Cell cycle analysis was performed by propidium iodide staining and flow cytometry. The NTC and KSR1 KO H82 cells were treated with 16 µM cisplatin (Sigma #232120) for 48 hours. Cisplatin was removed by washing four times with PBS, and fresh medium was replaced. The cells were harvested 0-72 h post-treatment along with untreated cells, fixed in cold 70% ethanol, and stained for DNA content using Telford reagent (Sigma #4864). DNA was analyzed by flow cytometry (FACSCalibur).

### Multi-well Resistance Assay

Multi-well resistance assays were performed and analyzed as recently described^(49,50)^. Cells were plated at low density (1000-2000 cells/well) in replicate 96-well plates (Fisher #237105), and each plate was treated with the indicated doses of cisplatin (Sigma #232120) or ERK inhibitor, SCH772984 (SelleckChem #S7101). Wells were fed with complete media and assessed weekly for outgrowth, wells that were >50% confluent were scored as resistant to the given dose of cisplatin. Data were plotted similar to a Kaplan-Meier curve, with cisplatin sensitivity (100 = no wells with >50% confluency) as the dependent variable (y-axis) and time in weeks as the independent variable (x-axis). Significance between curves was assessed using GraphPad Software, Inc.

### Sequencing Analysis

RNA sequencing data was obtained from 51 Patient-derived xenografts (PDXs)^(51)^. A correlation analysis was done between KSR1 and delta-AUC. Based on the biphasic clinical trajectory of SCLC, the delta-AUC is a measure of the drug response of PDXs to different chemotherapy regimens; cisplatin + etoposide (E+P), olaparib + temozolomde (O+T), and topotecan.

### Statistical Analysis

All western blots were performed a minimum of three times using independently generated cell lines to ensure the reproducibility. Statistical significance for colony forming assays was determined by Student’s t-test, to correct for independent groups’ comparison. For *in vitro and in vivo* ELDAs, statistical significance was determined by chi-squared test, since the assays deal with categorical variables in the sample. For dose-response curves, p values between NTC and KSR1 KO/ NTC + ERKi group were calculated by the extra sum-of-square F test. For growth curve and *in vivo* tumor volume analysis, statistical significance was determined by Student’s t-test for each day between the NTC and KSR1 KO group. For cell cycle analysis, the experiment was done three times, and all data are presented as the mean ± standard deviation. For multi-well resistance assays, significance was assessed by comparing Kaplan-Meyer Meier (survival analysis) curves for three independent experiments.

## Key Resources Table

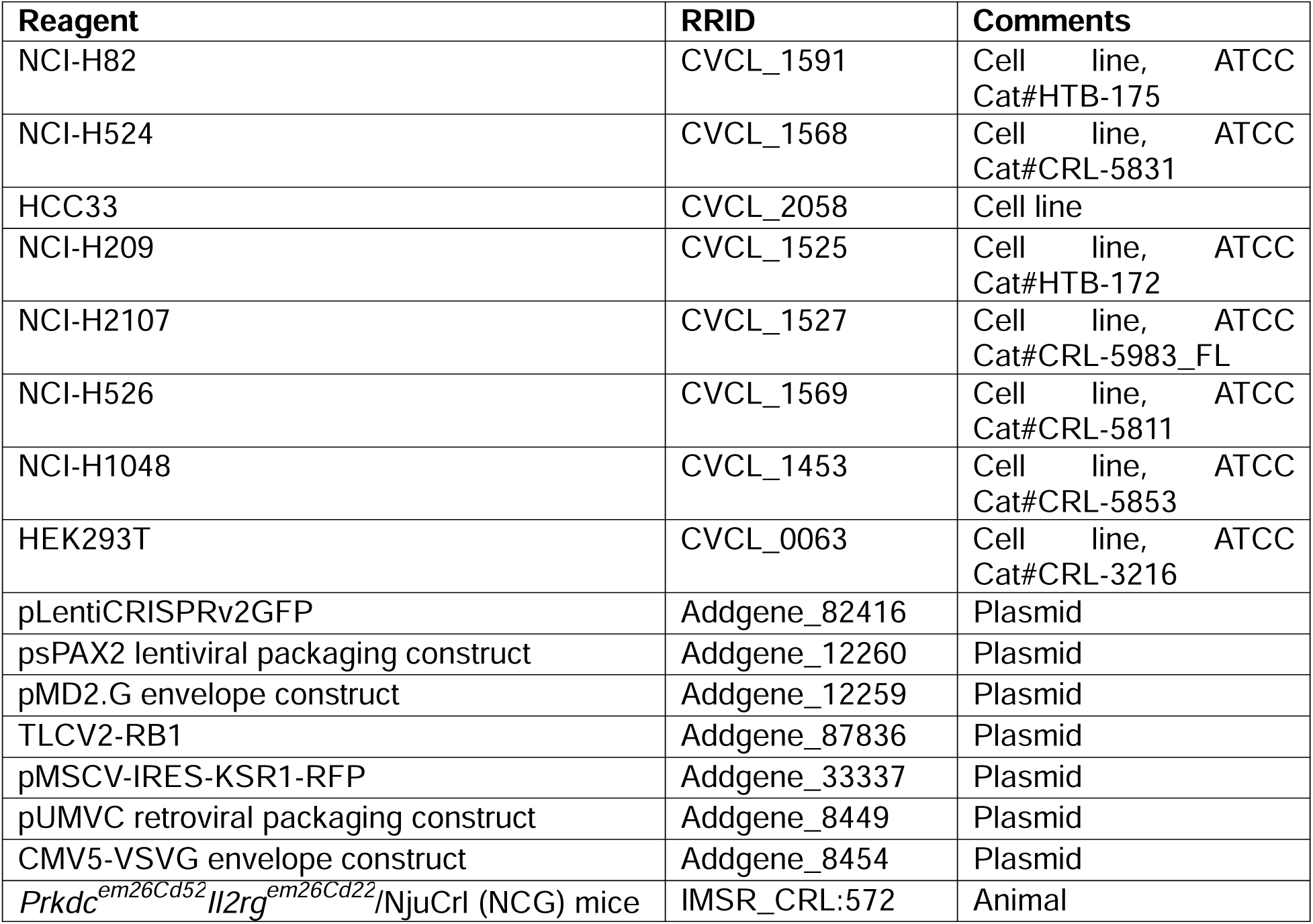
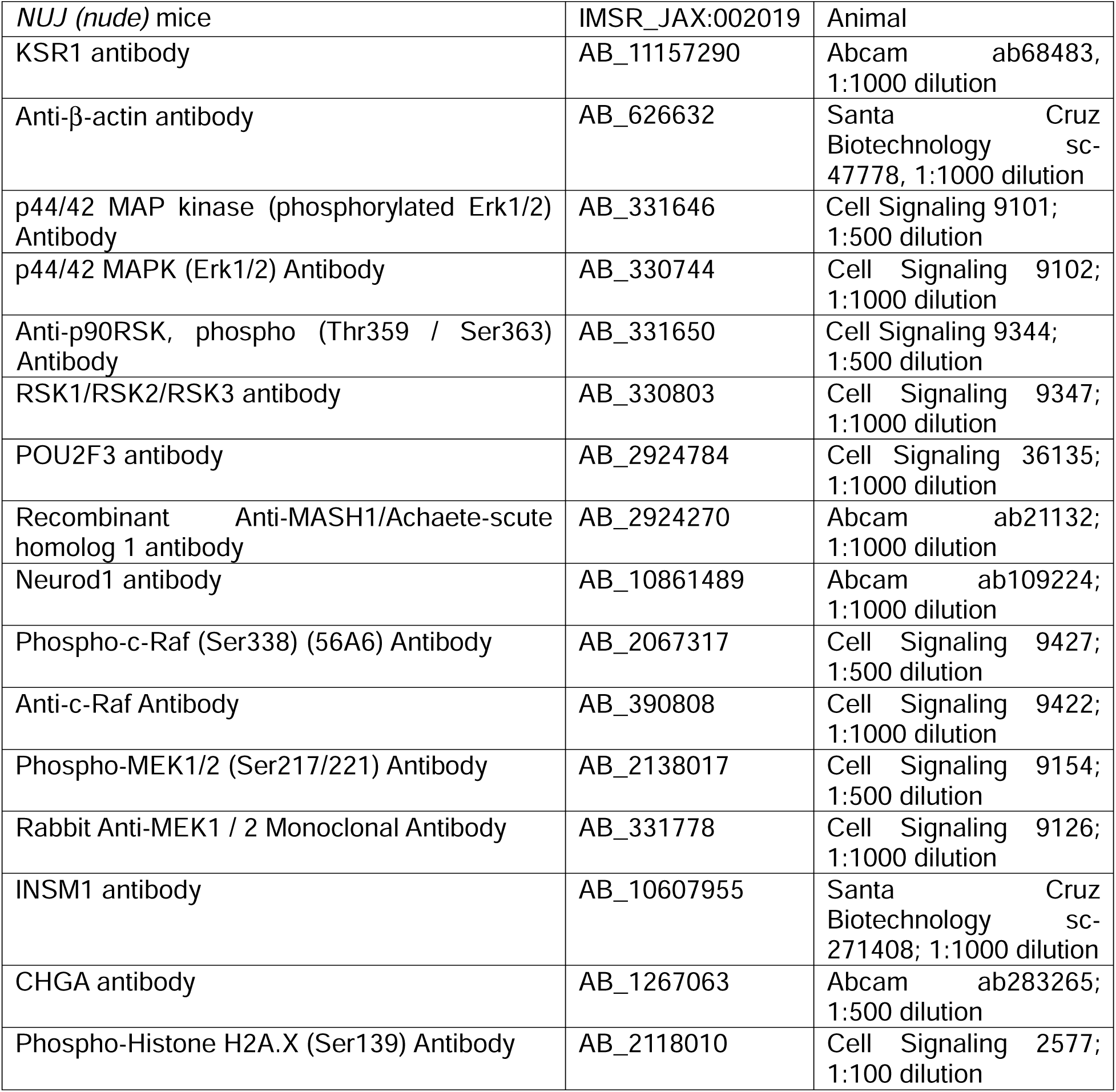

### KSR1 sgRNA sequences

sg1 5’ TTGGATGCGCGGCGGGAAAG 3’

sg2 5’ CTGACACGGAGATGGAGCGT 3’

NTC sgRNA sequence: 5’ CCATATCGGGGCGAGACATG 3’

### Data Availability Statement

The data generated in this study are available upon request from the corresponding author.

## Results

### KSR1 is required for SCLC cell clonogenicity

SCLC tumors harbor subpopulations of both clonogenic and self-renewing TICs and bulk non-TICs^(19,22)^. In contrast to the TICs, which are crucial for initial establishment, long term propagation, and metastatic ability of the tumors, the non-TIC population is highly proliferative but is inefficient at establishing new tumors *in vivo*^(19)^. In Ras-mutated colorectal and non-small cell lung cancer cell lines, depletion of KSR1 suppressed transformation and prevented drug-induced TIC upregulation^(46,47)^. Since KSR1 is expressed in multiple SCLC subtypes **<u>(Supplementary Fig. 1A)</u>** and KSR2 regulates self-renewal and clonogenicity of SCLC-A (submitted) (bioRxiv 2022.02.11.480157v4), we tested the ability of KSR1 disruption to impair clonogenicity and TIC formation in these cells. KSR1 was targeted in SCLC-N cell lines H82, H524, and HCC33 by constitutive or DOX-induced expression of Cas9 and one of two distinct sgRNAs (sg1, sg2) **(Fig. 1A, B, C, D <u>upper panels)</u>**. KSR1 was also targeted in SCLC-POU2F3 cell lines H526 and H1048 **(Fig. 1E**) and SCLC-A cell lines H209 and H2107 **(Fig. 1F**) by constitutive expression of Cas9 and one of two distinct sgRNAs (sg1, sg2). Clonogenicity is an *in vitro* index of self-renewal, a key property of TICs^(19)^. Clonogenicity was tested through single-cell colony forming assays^(19,49)^. Following confirmation of KSR1 knockout by western blotting, NTC and KSR1 KO cells were sorted by FACS and plated as single cells in 96-well plates to be analyzed for colony formation by CellTiter-Glo®. Robust colony formation was observed for the control SCLC-N/POU2F3 cells that was significantly suppressed upon KSR1 knockout in all five cell lines **(Fig. 1A, B, C, D, <u>lower panels, Fig. G, H)</u>**. These data demonstrate that KSR1 is a significant contributor to the clonogenic properties of SCLC TICs.

**Fig. 1.**
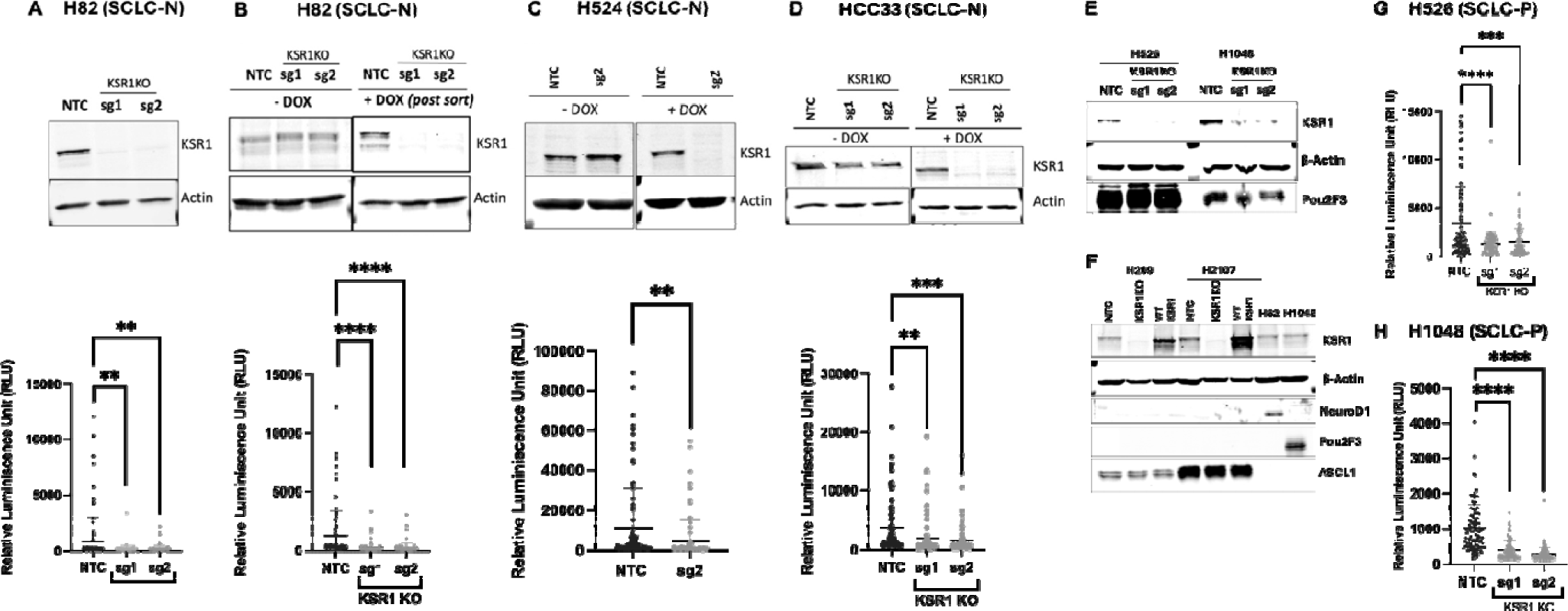
KSR1 is required for SCLC clonogenicity. **(A-D,** upper panels) KSR1 Western blot and (lower panels) clonogenicity of H82 cells following constitutive **(A)** or DOX-inducible **(B)** CRISPR/Cas9 in non-targeting control (NTC) or KSR1 KO (sg1, sg2 gRNAs) **(A, B)** H82, **(C)** H524 and **(D)** HCC33 cells. **(E, F)** KSR1, POU2F3 and **(F)** NEUROD1, ASCL1 Western blots from NTC and KSR1 KO **(E)** H526 and H1048, and **(F)** H209 and H2107 cells. **(G-H)** Clonogenicity of NTC and KSR1 KO **(G)** H526 and **(H)** H1048 cells. Blots are representative of three independent experiments. **, p<0.01; ****, p<0.0001.

### KSR1 disruption inhibits TIC frequency in SCLC cells

ELDA is a marker-free approach measuring TIC clonogenicity that estimates stem cell frequency from limiting dilution analysis^(48)^. ELDA, when performed *in vitro* determines the stem cell frequency within the bulk tumor cell population based on whether a spheroid is formed or not formed at a variety of cell dilutions. *In vitro* ELDA revealed a 3-5-fold dependency of TICs on KSR1 expression across NeuroD1, ASCL1 and POU2F3 SCLC subtypes. In the SCLC-N cell line H82, TIC frequency decreased from 1 TIC/3.36 tumor cells in NTC to 1 TIC/11 tumor cells in KSR1 KO cells **(Fig. 2A, <u>Supplementary Table 1</u>)**. Similarly, in the SCLC-POU2F3 cell line H1048, CRISPR targeting KSR1 decreased TIC frequency 5-fold from 1 TIC/14.4 tumor cells in NTC cells to 1 TIC/69.1 tumor cells in KSR1 KO cells **(Fig. 2E, <u>Supplementary Table 1)</u>**. This dependence upon KSR1 was replicated in SCLC-A cell line H209 where CRISPR targeting KSR1 decreased TIC frequency 3.5-fold from 1 TIC/30 tumor cells in NTC cells to 1 TIC/107.5 tumor cells in KSR1 KO cells **(Fig. 2F, <u>Supplementary Table 1)</u>**. Similar trends were observed for other SCLC-N cell lines H524 **(Fig. 2B, <u>Supplementary Table 1)</u>** and HCC33 **(Fig. 2C, <u>Supplementary Table 1)</u>**, SCLC-POU2F3 cell line H526 **(Fig. 2D, <u>Supplementary Table 1)</u>** and SCLC-A cell line H2107 **(Fig. 2G**). These data demonstrate a significant dependence of SCLC TICs on KSR1. The frequency of TICs is low in SCLC-A cells compared to SCLC-N or SCLC-POU2F3 cell lines. Since H209 and H2107 control cells only have 1TIC/30-40 tumor cells **(Fig. 2F, 2G, <u>Supplementary Table 1)</u>**, it is difficult to obtain colonies from 96 wells from single cell colony forming assays, hence clonogenicity assays could not be performed for SCLC-A cells.

**Fig. 2.**
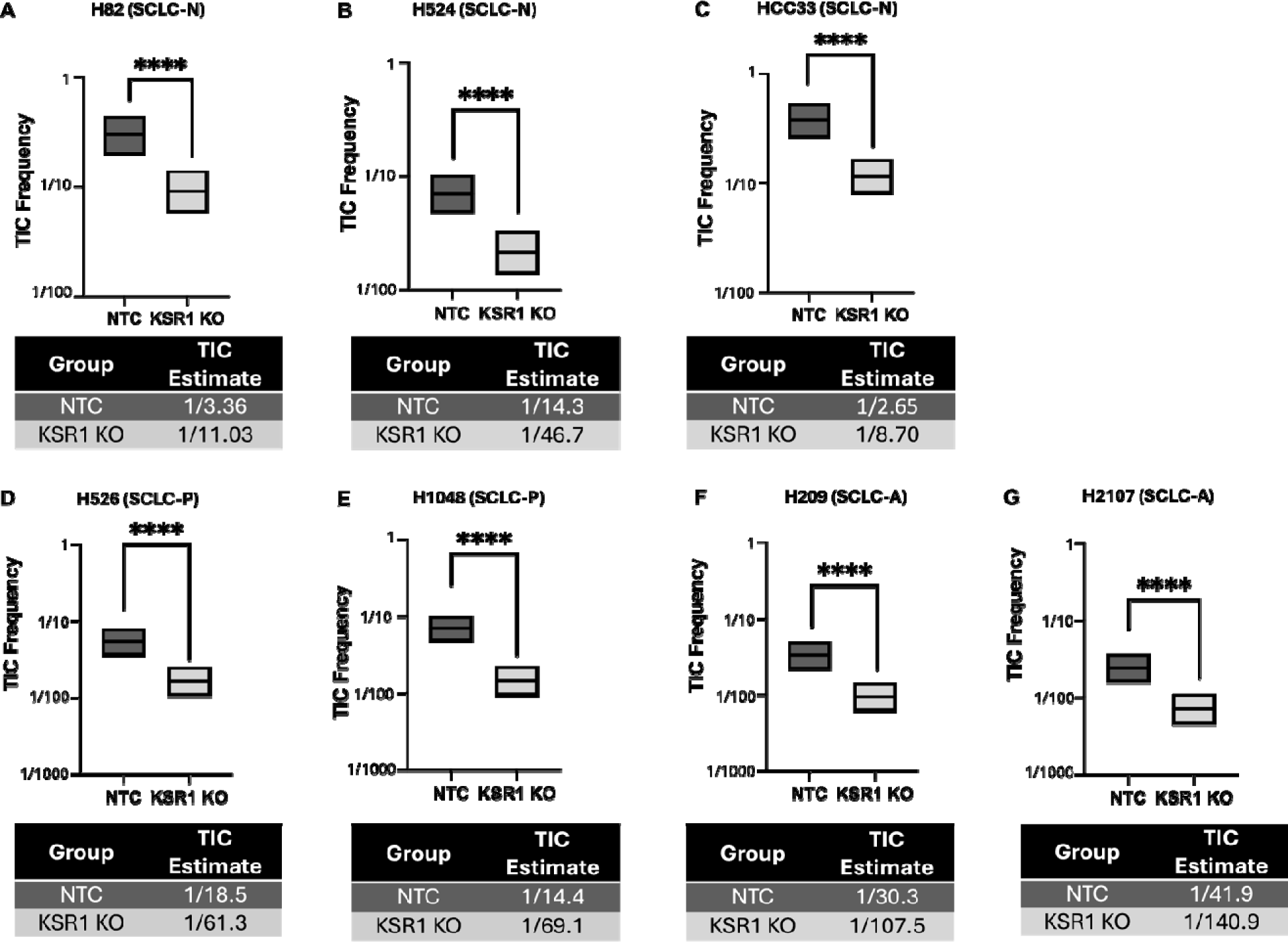
KSR1 is required for tumor initiation in NeuroD1, POU2F3, and ASCL1 subtype SCLC cell lines. *In vitro* ELDA of non-targeting control (NTC) and KSR1 KO **(A)** H82, **(B)** H524, **(C)** HCC33**, (D)** H526, **(E)** H1048, **(F)** H209, and **(G)** H2107 cells. Graphs are representative of three independent experiments. ****, p<0.0001.

### KSR1-dependent tumor initiation requires interaction with ERK

To test for off-target effects from CRISPR-Cas9 targeting a construct containing wildtype murine KSR1 (WTKSR1) that cannot be recognized by the sgRNAs (sg1, sg2) was expressed in KSR1 KO H82 cells **(Fig. 3A**). WTKSR1 expression restored colony formation to wildtype levels in these cells **(Fig. 3B**). *In vitro* ELDA assays reveal a restoration of the TIC frequency to 1 TIC/2.68 tumor cells in WTKSR1 cells, comparable to the TIC frequency of 1 TIC/3.66 tumor cells in control H82 cells **(Fig. 3C**). These data confirm the specificity of KSR1 CRISPR targeting.

**Fig. 3.**
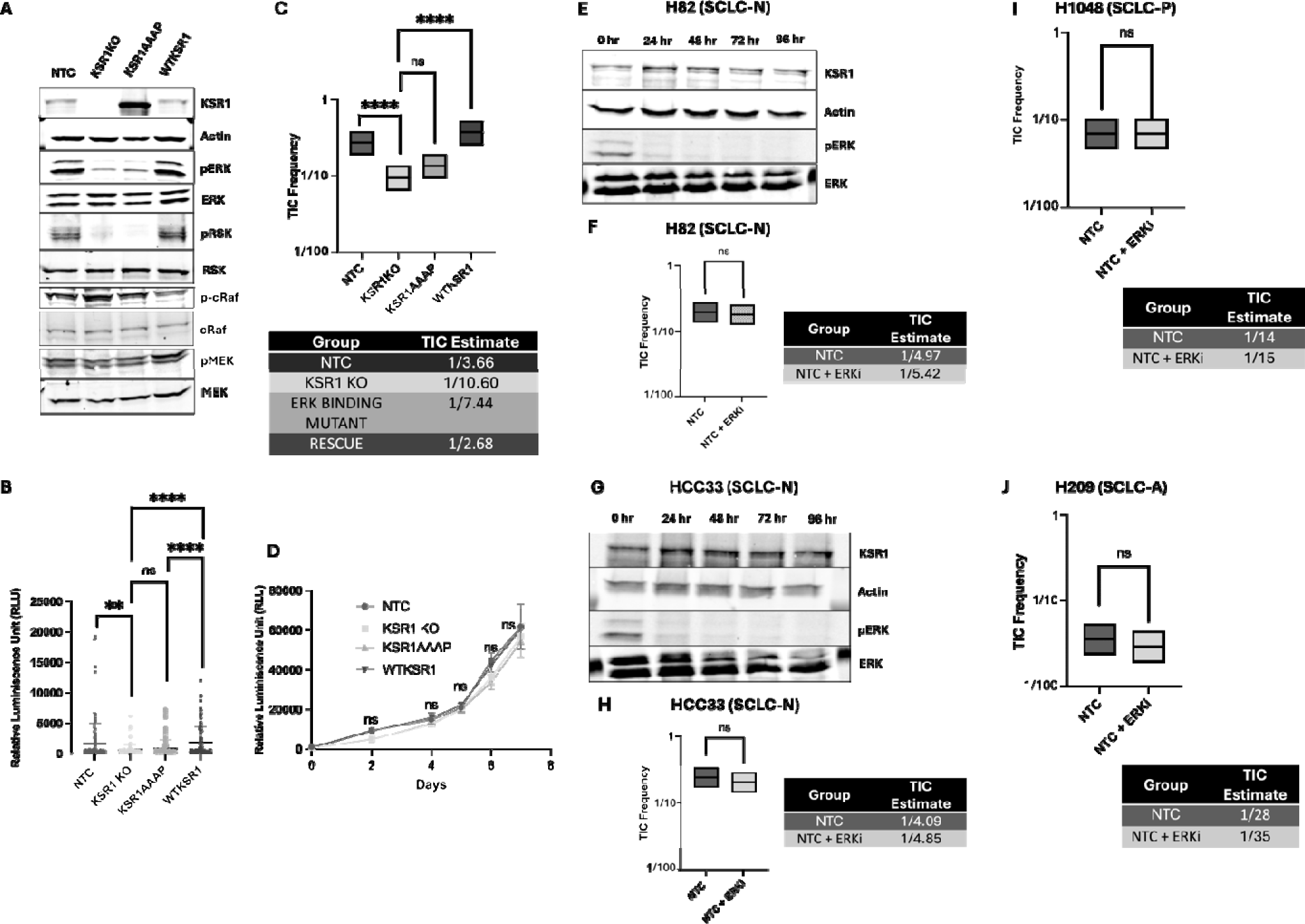
KSR1-ERK interaction but not ERK activity is required for H82 TIC formation. **(A)** Western blot of phospho- and total ERK, Rsk, Raf and MEK, H82 cells without (NTC) and with (KSR1 KO) DOX-inducible targeting of KSR1 and rescued with KSR1AAAP and WTKSR1. **(B)** Clonogenicity, **(C)** *in vitro* ELDA, and **(D)** Growth curves of NTC and KSR1 KO H82 cells without or with KSR1AAAP, or KSR1 cells. **(E, G)** Western blot of phospho- and total ERK, and **(F, H)** *in vitro* ELDA in NTC (**E, F)** H82 **(G, H),** HCC33, **(I)** H1048, and **(J)** H209 cells with 2 µM SCH772984 (ERKi). Results are representative of three independent experiments. ns, not significant; **, p<0.01; ****, p<0.0001.

The molecular scaffold KSR1 may contribute to TIC formation by promoting efficient signaling through the RAF/MEK/ERK kinase cascade^(37,38,41–44)^. To determine if KSR1-regulated ERK signaling pathways are important in SCLC-N tumor initiation, a construct encoding a DEF-domain (KSR1AAAP) mutated KSR1 transgene deficient in ERK binding^(37,42)^ was expressed in KSR1 KO H82 cells. Knockout of KSR1 reduces activation of ERK signaling as measured through phosphorylation of p90 ribosomal S6 kinase (RSK), key downstream substrates of ERK^(52)^ in the H82 cell line, that is rescued by the addition of the WTKSR1 transgene. The phosphorylation of substrates upstream of ERK, like c-Raf and MEK does not change in KSR1 KO and KSR1AAAP H82 cells **(Fig. 3A**). Unlike WTKSR1, the KSR1AAAP transgene is unable to restore *in vitro* colony formation to WT levels **(Fig. 3B**). *In vitro* ELDA assays reveal a TIC frequency of 1 TIC/7.44 tumor cells for the ERK binding mutant (KSR1AAAP) cells, similar to the TIC frequency observed for the *KSR1* KO cells, demonstrating that KSR1 incapable of interaction with ERK is also unable to restore the TIC frequency **(Fig. 3C, <u>Supplementary Table 1)</u>**. Growth curves, monitored over one week show that control, KSR1 KO, KSR1AAAP and WTKSR1 cells grow at similar rates **(Fig. 3D)**. These data show that KSR1-dependent ERK signaling is conserved in H82 cells, suggesting that ERK signaling may contribute to SCLC TIC formation.

To evaluate the contribution of ERK activity to SCLC TIC formation, *in vitro* ELDA assays were performed across all SCLC subtypes with non-targeting control (NTC) cells in the presence or absence of SCH772984, a novel, specific, ATP competitive inhibitor of ERK1/2 that exhibits robust efficacy in RAS- or BRAF-mutant cancer cells^(53–55)^. Treatment with 2 µM SCH772984 reduces phosphorylation of ERK over a period of 96 hours in SCLC-N cell lines, H82 **(Fig. 3E**) and HCC33 **(Fig. 3G**). However, *in vitro* ELDA assays performed with or without 2 µM SCH772984 fail to change the TIC frequency between the untreated and treated groups in H82 (SCLC-N) **(Fig. 3F, <u>Supplementary Table 1)</u>**, HCC33 (SCLC-N) **(Fig. 3H, <u>Supplementary Table 1)</u>**, H1048 (SCLC-POU2F3) **(Fig. 3I, <u>Supplementary Table 1),</u>** and H209 (SCLC-A) **(Fig. 3J, <u>Supplementary Table 1)</u>**, suggesting that ERK activity does not play a role in SCLC TIC formation.

### KSR1 disruption inhibits the tumor initiating capacity of SCLC-N cells *in vivo*

*In vivo* ELDA determines the stem cell frequency within the bulk tumor cell population from the frequency of tumor-positive and tumor-negative injections within a range of transplant doses. To test the effect of KSR1 disruption *in vivo*, control and KSR1 KO H82 cells were injected with successive dilutions subcutaneously into NCG mice. Tumors were monitored until the first tumor reached 1 cm^2^ and then all mice were sacrificed. The presence and absence of tumor formation was scored as “1” and “0” respectively. KSR1 disruption reduced frequency of TICs approximately 3-fold, from 1/62.5 control (NTC) H82 cells to 1/205.6 KSR1 KO H82 cells **(Fig. 4, <u>Supplementary Table 1)</u>**. These data support the conclusion from *in vitro* ELDA that KSR1 is an important effector of self-renewal, clonogenicity and tumor initiation in SCLC-N TICs.

**Fig. 4.**
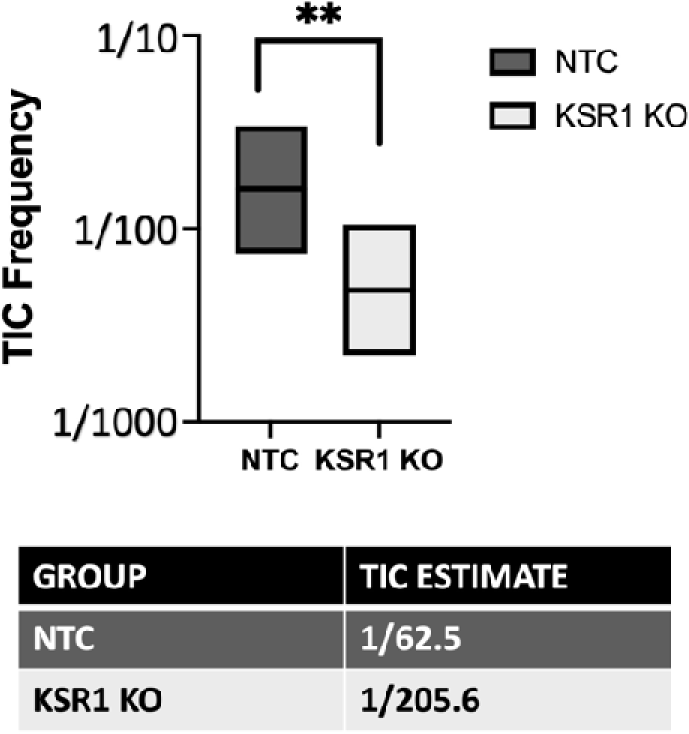
KSR1 knockout inhibits SCLC tumor initiation *in vivo*. The proportion of TICs in H82 cells was determined by *in vivo* ELDA of non-targeting control (NTC) and KSR1 KO cells. **, p<0.01, n=3 for each cell dilution.

### KSR1 disruption enhances cisplatin toxicity towards SCLC TICs

Given the contribution of KSR1 in promoting TIC formation and the role of TICs in therapeutic resistance^(1,2,5–9)^, we tested the effect of KSR1 disruption on SCLC response to the first line chemotherapeutic, cisplatin. To establish the doses of cisplatin that inhibit survival (≥ EC50) control and KSR1 KO NeuroD1 subtype cell lines H82, H524, and HCC33, ASCL1 subtype cell line H209, and POU2F3 subtype cell line H1048 were treated with increasing doses of cisplatin and cell viability was assessed 72 hours after treatment **<u>(Supplementary Fig. 3)</u>**. Dose-response curves revealed that in all three SCLC-N cells, KSR1 disruption modestly reduced the ED50. The doses of cisplatin that inhibited survival were higher in H82 than the other two SCLC-N cell lines. The SCLC-POU2F3 cell line, H1048 exhibited an ED50 similar to that of the H82 line, while H209 (SCLC-A) was more sensitive. To understand KSR1-dependent TIC formation in the context of cisplatin resistance, we performed *in vitro* ELDAs with the NTC and KSR1 KO SCLC cells with and without cisplatin. The TIC frequency in H82 cells decreases by 3-fold in the absence of any drug treatment in KSR1 KO (1 TIC/7 tumor cells) versus control (1 TIC/2 tumor cells) cells. ED85 cisplatin treatment reduced the TIC frequency in KSR1 KO cells by almost 7-fold; from 1 TIC/7 tumor cells in KSR1 KO cells without treatment to 1 TIC/46 tumor cells in KSR1 KO H82 cells with ED85 cisplatin treatment **(Fig. 5A, <u>Supplementary Table 1)</u>**. These data suggested no more than an additive effect for KSR1 KO in combination with cisplatin. For the more cisplatin-sensitive H524 and HCC33 cell lines, we tested cisplatin ED25, ED50, and ED75 doses in control and KSR1 KO cells via *in vitro* ELDA and observed a similar response to cisplatin combined with KSR1 KO. While there is an expected decrease in the TIC frequency of 3-4-fold with KSR1 disruption compared to controls with each cisplatin dose, the TIC frequency decreased in a dose-dependent manner with increasing doses of cisplatin **(Fig. 5B, C, <u>Supplementary Table 1)</u>**.

**Fig. 5.**
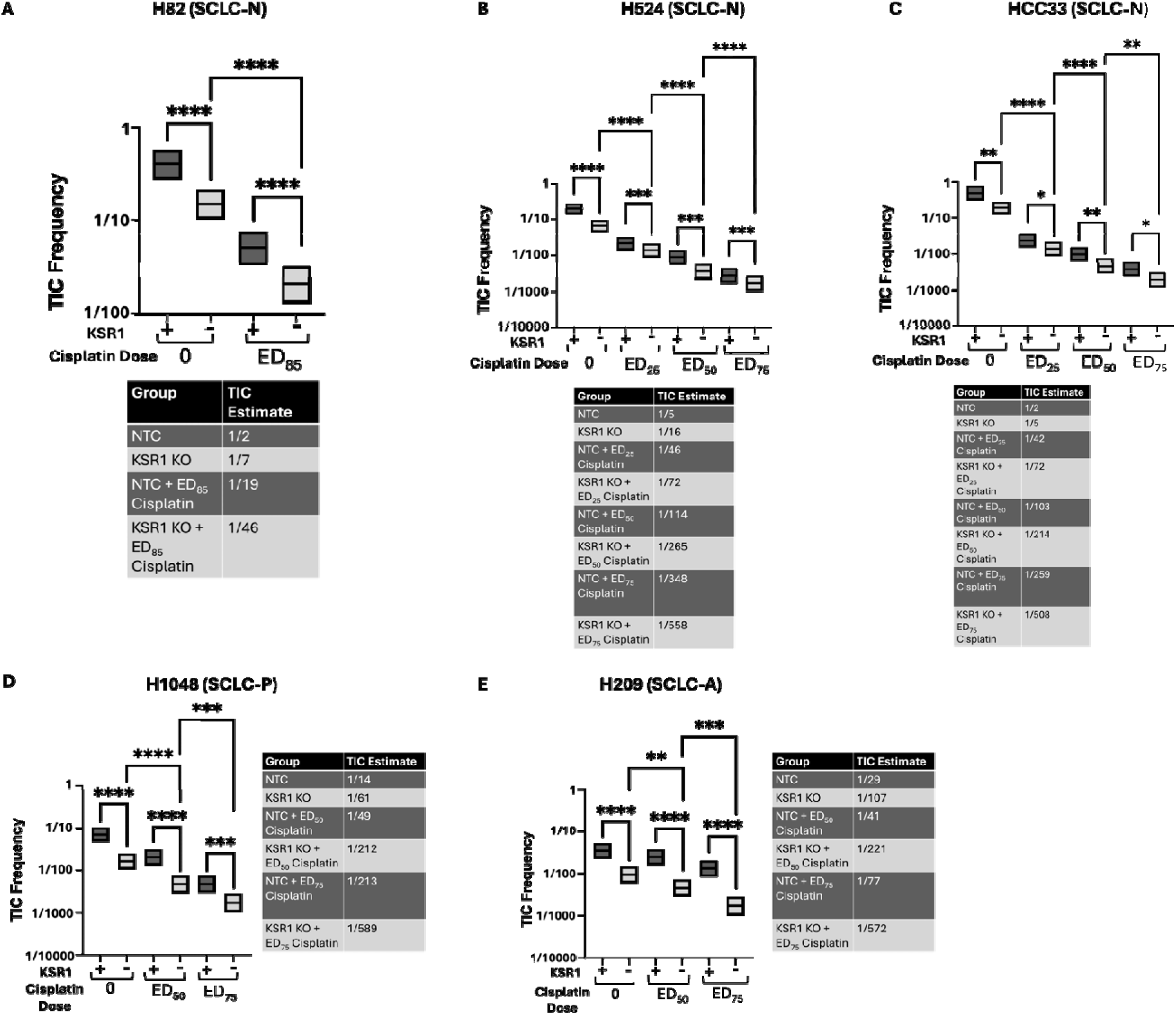
KSR1 promotes cisplatin resistance in NeuroD1, POU2F3, and ASCL1 subtype SCLC cell lines. *In vitro* ELDA of non-targeting control (NTC) and KSR1 KO **(A)** H82, **(B)** H524, **(C)** HCC33, **(D)** H1048, and **(E)** H209 cells using the indicated doses of cisplatin for 7-9 days. *, p<0.05; **, p<0.01; ***, p<0.001; ****, p<0.0001.

A similar trend was observed for H209 (SCLC-A) and H1048 (SCLC-POU2F3) cell lines. In POU2F3 subtype line H1048, while there is an expected decrease in the TIC frequency of 3-4-fold with KSR1 disruption as compared to controls with each cisplatin dose, the TIC frequency decreases by about 9-fold in KSR1 KO cells without treatment (1 TIC/61 tumor cells) vs KSR1 KO cells with ED75 treatment (1 TIC/589 tumor cells) **(Fig. 5D, <u>Supplementary Table 1)</u>**. In H209 (SCLC-A), there is a 3-fold decrease in the TIC frequency with KSR1 KO in the absence of any drug treatment. Cisplatin has a dose-dependent effect on TIC formation abundance in each cell line, but the effect of KSR KO is additive to each dose. The difference becomes 5-fold between the control (1 TIC/41 tumor cells) vs KSR1 KO (1 TIC/221 tumor cells) cells with ED50 cisplatin treatment and 7-fold between the control (1 TIC/77 tumor cells) vs KSR1 KO (1 TIC/572 tumor cells) cells with ED75 cisplatin treatment **(Fig. 5E, <u>Supplementary Table 1)</u>**. These data indicate a consistent decrease in the TIC frequency between control and KSR1 KO groups, with cisplatin treatment showing an additive effect by further decreasing TICs in a dose-dependent manner. These data highlight the role of KSR1 KO in enhancing cisplatin toxicity towards SCLC TICs, across all the different subtypes.

### KSR1 disruption prevents cisplatin resistance in SCLC cells

The ability of KSR1 KO to enhance cisplatin toxicity toward SCLC TICs **(Fig. 5**) and previous data indicating that TICs are synonymous with drug tolerant persister cells^(32,56,57)^, led us to test whether KSR1 disruption has any effect on SCLC cisplatin resistance. The similarity between control and KSR1 KO SCLC cell lines to cisplatin **<u>(Supplementary Fig. 3)</u>** allowed us to evaluate resistance using the same cisplatin dose in each control and cognate KO cell line in a multi-well resistance assay ^(49,50)^. Control and KSR1 KO cells were plated at low density (1000 cells/well; <10% confluent) in multiple 96-well plates and each plate was either left untreated (to assess the time required for normal cell outgrowth) or treated weekly with a single inhibitory dose of cisplatin (ED85/ED75) for up to 14-15 weeks. All 96 wells of each plate were scored weekly and wells that were ≥ 50% confluent were scored as resistant to that dose of cisplatin. An absence of resistant cells indicates long term drug sensitivity. Percent cisplatin sensitivity (growth inhibition) was plotted as the dependent variable on the y-axis versus time, similar to Kaplan-Meyer curves. KSR1 KO prevented the outgrowth of resistant colonies in all the SCLC cell lines, representing the different subtypes.

Multi-well resistance assays revealed that cisplatin sensitivity was maintained for 11-12 weeks in KSR1 KO H82, H526, and H1048 cells, while more than 50% of control wells developed resistance in 6-7 weeks **(Fig. 6A, D, E,)**. KSR1 KO cell lines H524 (SCLC-N) HCC33 (SCLC-N), H209 (SCLC-A) and H2107 (SCLC-A) cells, which had a lower ED50 for cisplatin **<u>(Supplementary Fig. 3)</u>**, maintained cisplatin sensitivity for 13 weeks in, while more than 50% of control SCLC-N wells, and >70% for the SCLC-A wells, developed resistance in 5-6 weeks **(Fig. 6B, C, F, G**). While untreated NTC and KSR1 KO HCC33 cells grow out within 2-3 weeks, there is dose-dependent difference in the development of cisplatin resistance of control and KSR1 KO wells that becomes more pronounced with increasing concentrations of the drug **<u>(Supplementary Fig. 5)</u>**. The ability of KSR1 to promote SCLC cisplatin resistance **(Fig. 6**) and TIC formation (**Figs. 2, 3C, 4, 5**) is consistent with the suggestion that drug tolerance and tumor initiation arise from the same subset of SCLC cells. However, on doing a correlation analysis with RNA sequencing data from 51 Patient-derived xenografts (PDXs)^(51)^, we detect no correlation between KSR1 levels in cell lines or in patient-derived xenografts and their response to cisplatin + etoposide, olaparib + temozolomde, and topotecan **<u>(Supplementary Fig. 4)</u>**.

**Fig. 6.**
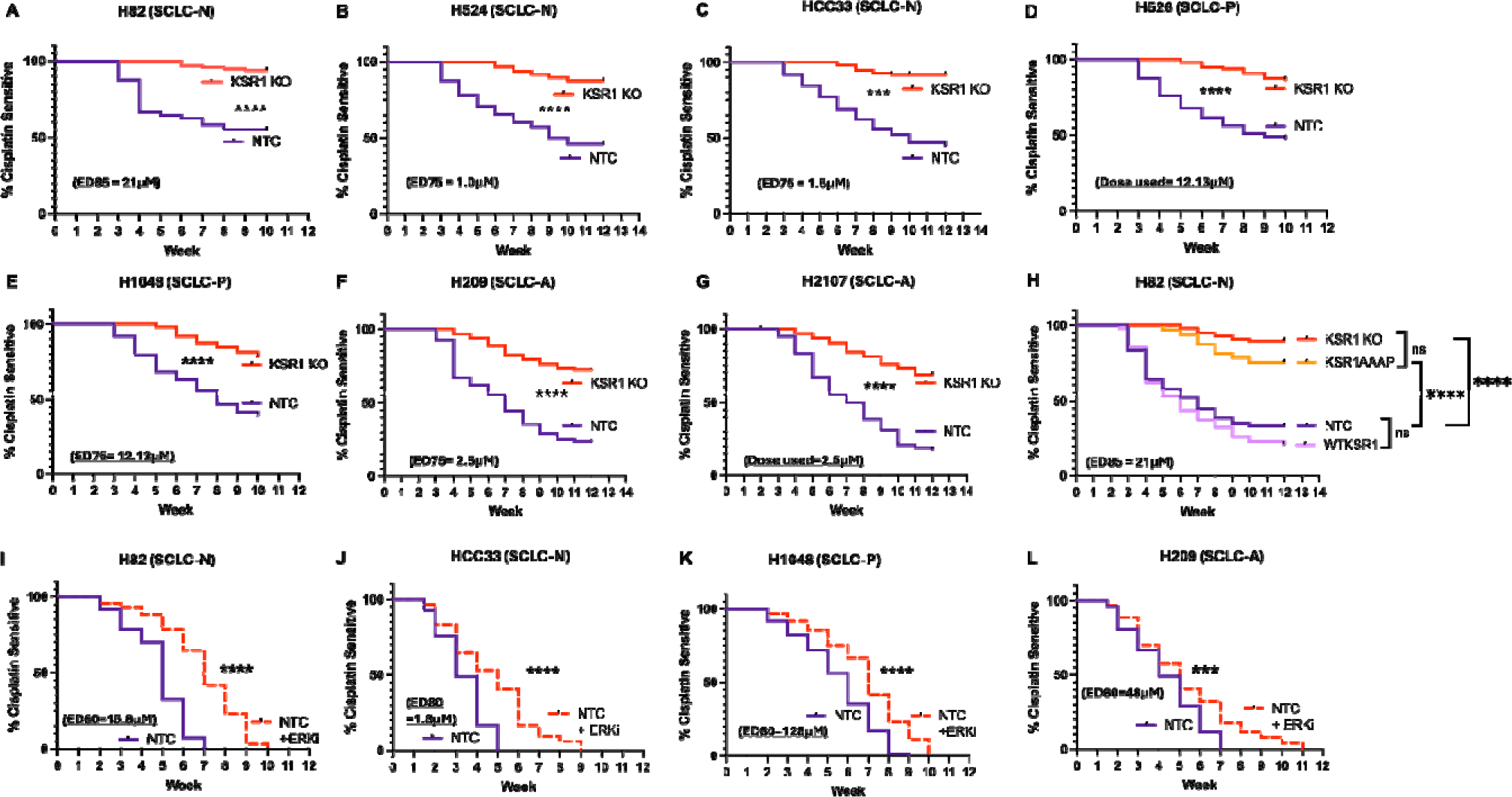
KSR1 knockout sensitizes SCLC cells to cisplatin. Multi-well resistance assay of NTC and KSR1 KO) **(A)** H82, **(B)** H524, **(C)** HCC33, **(D)** H526, **(E)** H1048, **(F)** H209, and **(G)** H2107 cells treated continuously with the indicated concentrations of cisplatin. **(H)** Multi-well resistance assay of NTC, KSR1 KO, KSR1AAAP and WTKSR1 H82 cells. **(I-L)** Multi-well resistance assay of NTC and NTC cells treated with 2 µM SCH772984 (ERKi) (NTC+ERKi) H82 **(I)**, HCC33 **(J)**, H1048 **(K)** and H209 **(L)** cells treated continuously with the indicated ED80 concentrations of cisplatin and scored weekly. One representative assay from experiments completed in triplicate is shown. ns, not significant; ***, p<0.001; ****, p<0.0001

Though the KSR1 DEF domain, which promotes interaction with ERK, is critical to TIC formation^(37,42,58)^, ERK kinase activity is dispensable (**Fig**. **3**). We examined the extent to which the interaction of KSR1 with ERK and ERK activity also contribute to cisplatin resistance. Multi-well resistance assays were performed with control H82 and KSR1 KO H82 cells, KSR1 KO H82 cells expressing a KSR1 transgene (WT KSR1), or the KSR1 construct unable to bind ERK (KSR1AAAP). While more than 70% control and rescue wells showed complete resistance in 8-9 weeks, only 10% of KSR1 KO wells and 25% of wells with KSR1 KO expressing the ERK binding mutant became cisplatin resistant **(Fig. 6D**). These data indicate that the ability of KSR1-dependent signaling to promote cisplatin resistance is dependent upon interaction with ERK.

To determine if MAPK pathway inhibitors could also prevent cisplatin resistance, dose-response curves were performed across multiple SCLC subtypes with non-targeting control (NTC) cells in the presence or absence of the MEK inhibitor Trametinib and the ERK inhibitor, SCH772984. The dose-response curves demonstrate no difference in the ED50 values between the drug-treated and untreated group for both Trametinib and SCH772984 **<u>(Supplementary Fig. 2)</u>**. Multi-well resistance assays demonstrate the ability of SCH772984 to modestly but significantly delay cisplatin resistance in control SCLC cells **(Fig. 6I-L**). However, unlike KSR1 KO **(Fig. 6A-H**), the ED80 dose of SCH772984 fails to prevent cisplatin resistance in any of the SCLC cell lines within the 3-month duration of the experiment. These data suggest that, while ERK activity makes a small contribution, the ability of KSR1 to interact with ERK is the predominant determinant to SCLC cisplatin resistance.

### KSR1 disruption sensitizes H82 tumors to cisplatin

The ability of KSR1 KO to sensitize SCLC cell lines to cisplatin in culture led us to test whether KSR1 disruption sensitizes SCLC tumor formation to cisplatin. 2.5 million non-targeting control (NTC) or KSR1-targeted (KSR1 KO) H82 cells were injected subcutaneously into immunocompromised mice. Six days after cell injection, mice with NTC or KSR1 KO tumors were divided into vehicle (PBS) and cisplatin treatment. Xenografted tumor growth was measured and plotted as tumor volume over time. Ten days after injection, vehicle-treated control and KSR1 KO cells formed palpable tumors that increased in volume at the same rate until sacrifice at day 20. Palpable tumors from control cells H82 cells were detectable in cisplatin treated mice at 18-20 days. Cisplatin-treated control tumors grew much slower than the tumors formed from vehicle treated control cells, eventually reaching 1000 mm^3^ at around 35 days. However, none of the cisplatin-treated mice injected with KSR1 KO H82 cells formed palpable tumors even after 35 days, demonstrating that KSR1 KO sensitizes the H82 tumor xenografts to cisplatin (**Fig. 7A**).

**Fig. 7.**
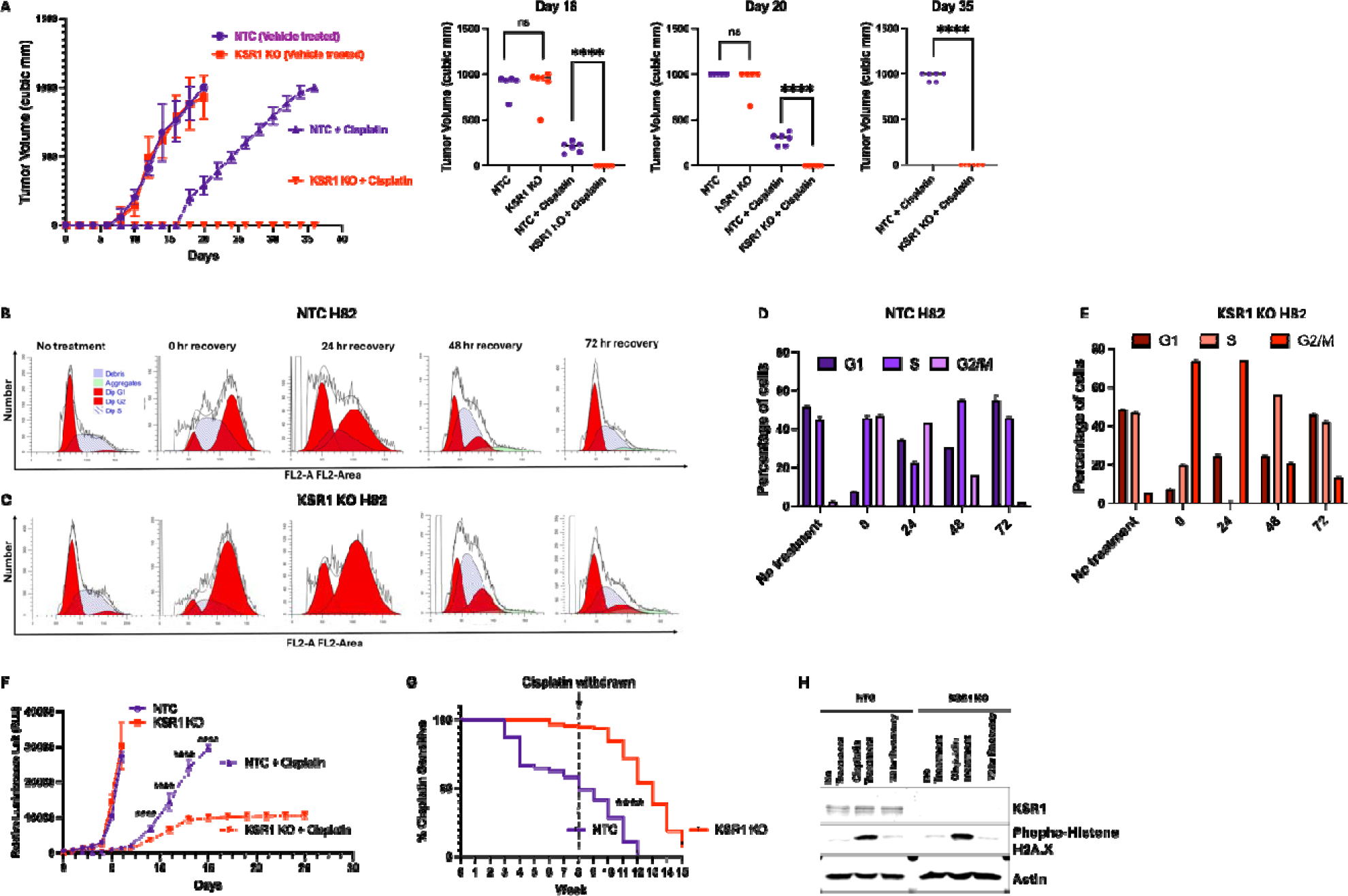
KSR1 knockout sensitizes H82 cells to cisplatin *in vivo and* disrupts G2/M. **(A)** Tumor volume of non-targeting control (NTC) and KSR1-targeted (KSR1 KO) H82 cells injected subcutaneously into nude mice with or without cisplatin treatment (n=6/group). **(B, C)** Cell cycle profiles and **(D, E)** quantification in non-targeting control (NTC), and KSR1 KO H82 cells without (No treatment) or with 48 hours of 16 µM (ED80) cisplatin. Cells were cleared of the drug and collected at 0, 24, 48, and 72 hours after cisplatin withdrawal. Data are presented as the mean ± standard deviation (n=3). **(F)** Growth curves of NTC and KSR1 KO H82 cells in the presence or absence of 16µM (ED80) cisplatin treatment. **(G)** Multi-well resistance assay of NTC and KSR1 KO H82 cells treated continuously with 16 µM cisplatin for 8 weeks and then withdrawn and scored for an additional 7 weeks. **(H)** Western blot of KSR1 and phospho-Histone H2A.X in control (NTC) and KSR1 KO H82 cells treated with and without 2 µM cisplatin for 48 hour, and 72 hours of recovery after withdrawal. ns, not significant; ****, p<0.0001

### KSR1 KO mediated cisplatin sensitivity in SCLC is not a function of cell cycle arrest or impaired cell growth

KSR1 is essential for reinitiation of the cell following DNA intra-strand crosslinking by mitomycin C^(58)^. To test the role of KSR1 in cell cycle re-entry following cisplatin induced DNA damage and its contribution to cisplatin resistance, we performed cell cycle analysis of cisplatin-treated cells by flow cytometry. Control and KSR1 KO H82 cells were treated with 16 µM cisplatin for 48 hours, which was necessary to cause cell cycle arrest. Cells were collected and stained with propidium iodide for flow cytometric analysis 0-72 hours after cisplatin removal. Compared to untreated cells (<5%), 48 hours of cisplatin treatment increased the proportion of G_2_/M control (45%) and KSR1 KO (72%) H82 cells lines indicating cell cycle arrest at the G_2_/M interface (**Fig. 7B, C, D, E**). KSR1-expressing control cells reinitiated the cell cycle, decreasing their G_2_/M population and increasing their G_1_ population within 24 hours after cisplatin withdrawal. Relative to control cells, release from G_2_/M arrest was delayed in the KSR1 KO cells, but most KO cells reentered the cell cycle 48 after cisplatin withdrawal. These data suggest both the control and KSR1 KO cells can recover and resume normal cell cycle following arrest, though KSR1 KO cell reentry is delayed.

Following DNA damage, histone H2AX is phosphorylated by ATM/ATR and localizes at sites of DNA double strand breaks (DSBs) until repair is complete^(59,60)^. 2 μM cisplatin treatment for 48 h causes a spike in the expression of phosphorylated histone H2AX (γ-H2AX) in control and KSR1 KO H82 cells, which returns to almost undetectable levels 72 hours after cisplatin withdrawal (**Fig. 7H**). These data demonstrate that the DNA damage does not persist in KSR1 KO SCLC cells.

Untreated control and KSR1 KO grow at similar rates. Cisplatin treated control cells grow much slower, but eventually the cisplatin treated control cells grow out to reach confluency comparable to the untreated ones implying that cisplatin significant delays but does not prevent the growth of control cell lines. Consistent with cisplatin-induced cell cycle delay (**Fig. 7C**), cisplatin-treated KSR1 KO cells grow, albeit at a slower rate than cisplatin-treated control cells, but never reach the same confluency as the cisplatin treated control cells even after 24 days, suggesting the cisplatin treated control and KSR1 KO cells plateau at different densities. (**Fig. 7F**).

To test whether SCLC cells recover from cisplatin-induced arrest in multi-well resistance assays, control and KSR1 KO H82 cells treated continuously with ED85 cisplatin for eight weeks, after which cisplatin was withdrawn from the assay. Although more than 90% of KSR1 KO cells remain sensitive to cisplatin after eight weeks of treatment, drug withdrawal allows KSR1 KO H82 cells to resume proliferation at rate comparable to control H82 cells. (**Fig. 7G**). These data indicate that KSR1-dependent cisplatin resistance in SCLC cells is not a function of an impaired cell cycle or abnormal cell growth.

## Discussion

Our data reveal an unexpected role of KSR1 in promoting tumor initiation and resistance to the first line chemotherapeutic cisplatin in SCLC cells. These data suggest that KSR1 increases the number of tumor initiating and drug tolerant cells in SCLC. Despite its ability to function as a Raf/MEK/ERK scaffold, the ability of KSR1 to enhance tumor initiation and cisplatin resistance appears dependent upon its interaction with ERK more than its ability to increase ERK activity.

Extensive genomic and molecular analysis led to classification of SCLC subtypes based on the differential expression of four key transcription factors, ASCL1, NeuroD1, POU2F3 and Yap1^(10,14)^, which has been recently reclassified as undifferentiated SMARCA4-deficient malignancies^(15)^. A fifth subtype based on the bHLH proneural transcription factor ATOH1, has also now been proposed^(16,17)^. Despite such detailed molecular characterization, all SCLC subtypes are treated with the same standard of care despite recent evidence that Myc-driven NeuroD1 tumors are responsive to therapies distinct from those effective against ASCL1 subtype tumors^(20,61)^. Our data demonstrate that KSR1 disruption impairs TIC formation and promotes cisplatin sensitivity in ASCL1, NeuroD1, and POU2F3 subtypes.

Tumor initiating cells (TICs) are a slow-growing subpopulation essential to tumor formation. TICs also serve as the reservoir of drug-tolerant persister (DTP) cells ^(25–31)^. Therefore, killing TICs, not just bulk tumor, is a key to limiting tumor dissemination and improving SCLC therapy. Isolating TICs is a key step in identifying their vulnerabilities. In ASCL1 subtype cell lines, TICs are enriched in CD24^high^CD44^low^ EPCAM^high^ subpopulations^(19)^ (bioRxiv 2022.02.11.480157v4). Manipulation of KSR1 in SCLC cells and tumors may facilitate the characterization of TICs and reveal their functional relationship, or identity, with DTPs. AT2+ bronchial epithelial cells and pulmonary neuroendocrine cells are cells-of-origin for SCLC^(18,62,63)^. The ability of KSR1 KO to suppress TIC formation in multiple subtypes that have different cells-of-origin suggests that common mechanisms underlie TIC formation regardless of the precursor or mutational spectrum.

The standard first-line chemotherapy regimen for limited stage SCLC is cisplatin–etoposide, which has not changed for the past three decades^(64)^. Patients with metastatic disease are treated with systemic chemotherapy with or without immunotherapy. Even though SCLC is exceptionally responsive to initial cytotoxic therapies, the majority of these responses are transient and exhibit a high incidence of relapse, in which case second-line chemotherapy is, in general, much less effective than initial treatment^(64,65)^. Inevitable relapse results in a median survival duration of approximately one year for patients with metastatic disease^(64)^. DTPs, a subpopulation of drug-tolerant cells found in tumors, play a critical role in the development of drug resistance. Like TICs, DTPs are slow cycling or dormant and maintain viability under therapy, suggesting a shared identity with TICs^(32)^. We detect no correlation between KSR1 levels in cell lines or in patient-derived xenografts and their response to cisplatin + etoposide, olaparib + temozolomde, and topotecan **<u>(Supplementary Fig. 4)</u>**. KSR1 KO has a modest effect on the dose-response of SCLC subtypes to cisplatin. KSR1-dependent cisplatin resistance is not predicted by the dose-dependent response of cell lines to cisplatin. Cisplatin ED50 of different SCLC cell lines does not correlate with the clinical history of the patient from which these cell lines were derived. SCLC-N cell lines H82, and H524 were derived from relapsed SCLC patients^(66)^ and have markedly different ED50s, whereas the SCLC-A cell line H209, which originated from a patient with treatment-naïve SCLC^(51)^, has a relatively intermediate ED50 for cisplatin. Estimated TIC frequency and SCLC subtype are also not predictors of KSR1-dependent cisplatin sensitivity. These data demonstrate that acute dose-response comparisons cannot capture or evaluate the emergence of long-term acquired cross-resistance, the major impediment to SCLC therapy^(51)^.

By combining elements of extended proliferation outgrowth ^(68,69)^ and time-to-progression assays ^(70)^, a novel multi-well resistance assay was established^(50)^. This assay models the development of acquired resistance over a 6- to 16-week time frame and has successfully modeled acquired resistance to several EGFR-RAS pathway targeted therapeutics including osimertinib, the KRAS^G12C^ inhibitors adagrasib and sotorasib, and the MEK inhibitor trametinib in multiple cancer cell lines^(49)^. Using this method, our study shows that KSR1 disruption significantly sensitizes SCLC cells to the first-line chemotherapy drug, cisplatin. The NeuroD1, ASCL1, and POU2F3 lines were similarly sensitized by KSR1 KO despite different predicted TIC frequencies (**Fig. 2**). These observations suggest that, like its effect on TIC formation, KSR1-dependent mechanisms promoting cisplatin resistance are intrinsic to multiple subtypes. The effects seen in resistance assays were replicated in tumor xenograft studies where tumors arose in engrafted and cisplatin-treated control H82 cells in less than three weeks but were undetectable in cisplatin-treated mice bearing KSR1 KO for term of the experiment (> 5 weeks).

SCLC tumors have loss-of function mutations in *p53* and *RB*^(14,67,68)^. These tumor suppressor genes have overlapping and distinct functions that facilitate cell cycle arrest or apoptosis in response to DNA damage^(69)^. We observed previously that KSR1 was required for cell cycle reentry following DNA damage and G_2_/M arrest in immortalized mouse embryo fibroblasts^(58)^. Despite impairment of these cell cycle checkpoints, H82 cells undergo G_2_/M cell cycle arrest following cisplatin treatment. However, both control and KSR1 KO cells recover and resume proliferation, albeit with a delayed reentry in KSR1 KO H82 cells **(Fig. 7E**). Consistent with cisplatin-induced cell cycle delay, cisplatin-treated KSR1 KO cells grow slower and reach a lower confluency compared to cisplatin-treated control cells, plateauing at different densities after 24 days **(Fig. 7F**). The reduced confluency of cisplatin-treated KSR1 KO cells may indicate that KSR1 KO cells, especially TICs, are more vulnerable to, and accumulate increasing amounts of unrepaired damage from cisplatin treatment that impair their ability to divide. Further investigations are needed to elucidate the precise KSR1-dependent mechanisms controlling cisplatin resistance, which could impact processes ranging from DNA repair to drug efflux.

KSR1 is a RAF/MEK/ERK scaffold, acting downstream of Ras and growth factor receptor tyrosine kinases to promote extracellular regulated kinase (ERK) signaling^(35,36)^. KSR1 disruption in Ras mutated colorectal and non-small cell lung cancer cell lines suppresses transformation and prevents TIC upregulation by the MEK inhibitor trametinib^(46,47)^. Activating Ras mutations are rare in SCLC, ERK activation is reported to induce cell-cycle arrest and senescence in ASCL1 subtype SCLC^(70,71)^, and low levels of phospho-ERK may be predictive of SCLC sensitivity to IGF1R inhibitors.^(72–75)^ Increased Raf/MEK/ERK signaling induced by expression of an activated Raf construct has been reported to cause growth arrest and reduce expression of neuroendocrine markers in SCLC^(74,76)^. Consistent with these observations, the ERK inhibitor SCH772984 has no effect on SCLC TIC formation and proliferation and, relative to KSR1 KO, a modest effect on cisplatin sensitivity. While ERK activity is not critical to the formation of SCLC TICs and DTPs, disruption of the KSR1 DEF domain, which is necessary for its interaction with ERK^(42,77,78)^, is critical to SCLC tumor initiation and cisplatin resistance. This raises the possibility that ERK is an allosteric modulator of KSR1 or that DEF domain mutations disrupt additional KSR1-mediated signals unappreciated for their contribution to TIC and DTP formation other than those conveyed by the role of KSR1 as an ERK scaffold^(45,79–81)^. Future work should be directed to test additional effectors of KSR1-dependent signaling on TIC formation and therapy resistance in SCLC.

The profound effect of KSR1 disruption on tumor initiation and cisplatin resistance across SCLC subtypes and its ability to acutely prevent cisplatin resistance SCLC tumor growth shows that KSR1 is a novel regulator of SCLC TIC clonogenicity both *in vitro* and *in vivo*, and a potential therapeutic target. Preclinical manipulation of KSR1 and identification of relevant downstream effectors may yield novel translatable approaches for SCLC therapy, which has shown only modest improvement in the last 40 years.^(1)^.

## Acknowledgements

The authors thank the UNMC Flow Cytometry Research Facility for help isolating the GFP+ SCLC cells. We declare no conflicts of interest.

## Author Contributions

D.C., D.H.H., and R.E.L. designed research; D.C., R.A.S., D.H.H., B.J.D., H.M.V., C.R., and J.W.A. performed research; D.C., D.H.H., B.J.D. and H.M.V. analyzed data; K.W.F., and R.E.L. edited manuscript; D.C. and R.E.L. wrote the paper.

