## Supplemental data for "KSR1 mediates small-cell lung carcinoma tumor initiation and cisplatin resistance"

**Supplementary Fig. 1.**

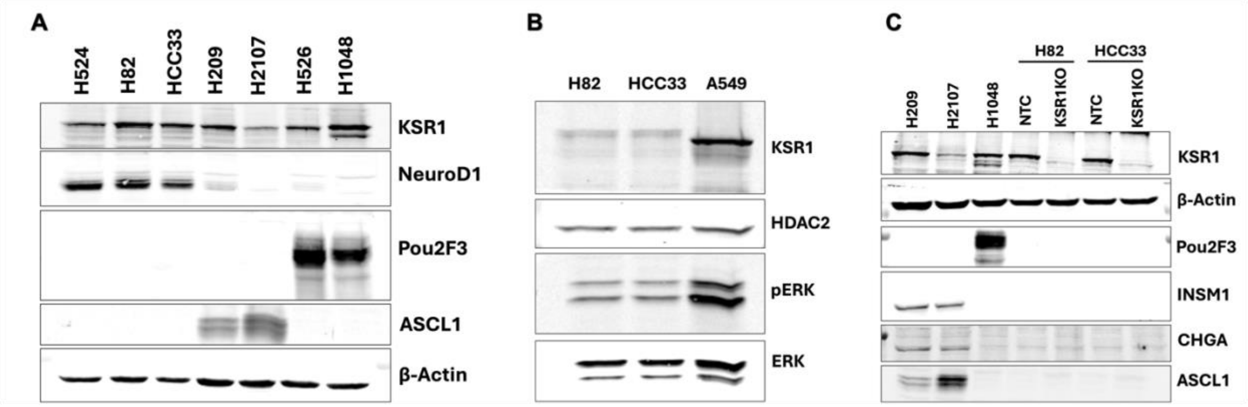


**Supplementary Fig. 1. Expression of key transcription factors and neuroendocrine markers across SCLC subtypes. (A)** Western blot of KSR1 and SCLC subtype markers NEUROD1, POU2F3 and ASCL1 in H524, H82, HCC33, H209, H2107, H526 and H1048 cells. **(B)** Western blot of phosphorylated (pERK) and total ERK in SCLC-N H82 and HCC33, and NSCLC A549 cells. **(C)** Western blot of KSR1, POU2F3, ASCL1 and neuroendocrine markers, INSM1 and CHGA levels in SCLC-A H209, H2017, SCLC-POU2F3 H1048, and control (NTC) and KSR1 KO SCLC-N H82 and HCC33 cells. Blots are representative of three independent experiments.

**Supplementary Fig. 2.**

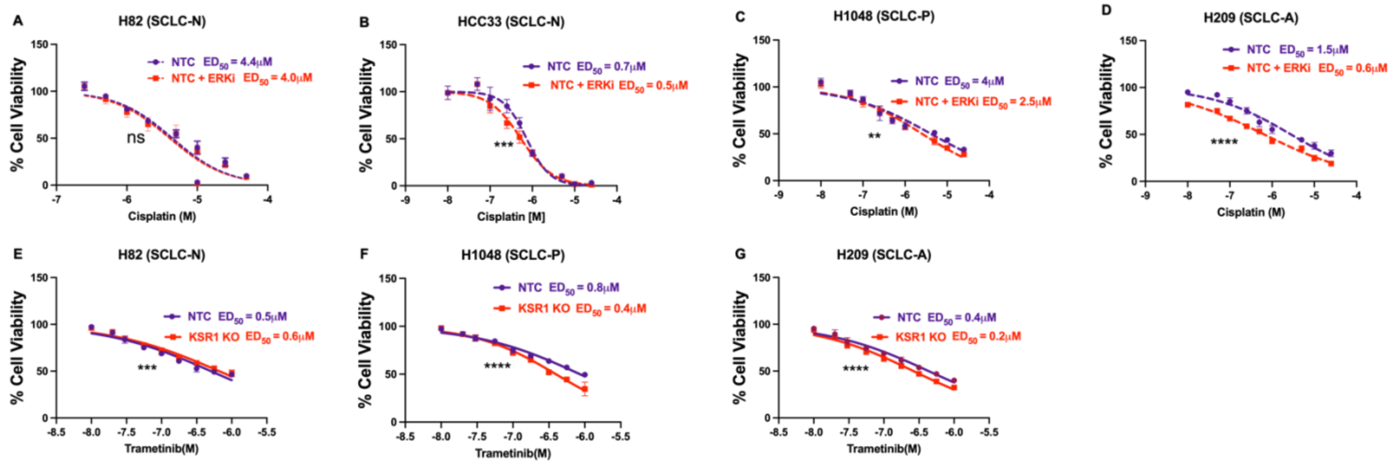


**Supplementary Fig. 2. SCLC cell lines show no to minimal sensitivity to MAPK pathway inhibitors. (A-D)** Cisplatin dose-response curves in non-targeting control **(A)** H82, **(B)** HCC33, **(C)** H1048, and **(D)** H209 cells, with and without 2 μM ERKi SCH772984. **(E-G)** Trametinib dose-response curve with non-targeting control (NTC), and KSR1 KO **(E)** H82, **(F)** H1048, and **(G)** H209 cells. ns, not significant; **, p<0.01; ***, p<0.001; ****, p<0.0001.

**Supplementary Fig. 3.**

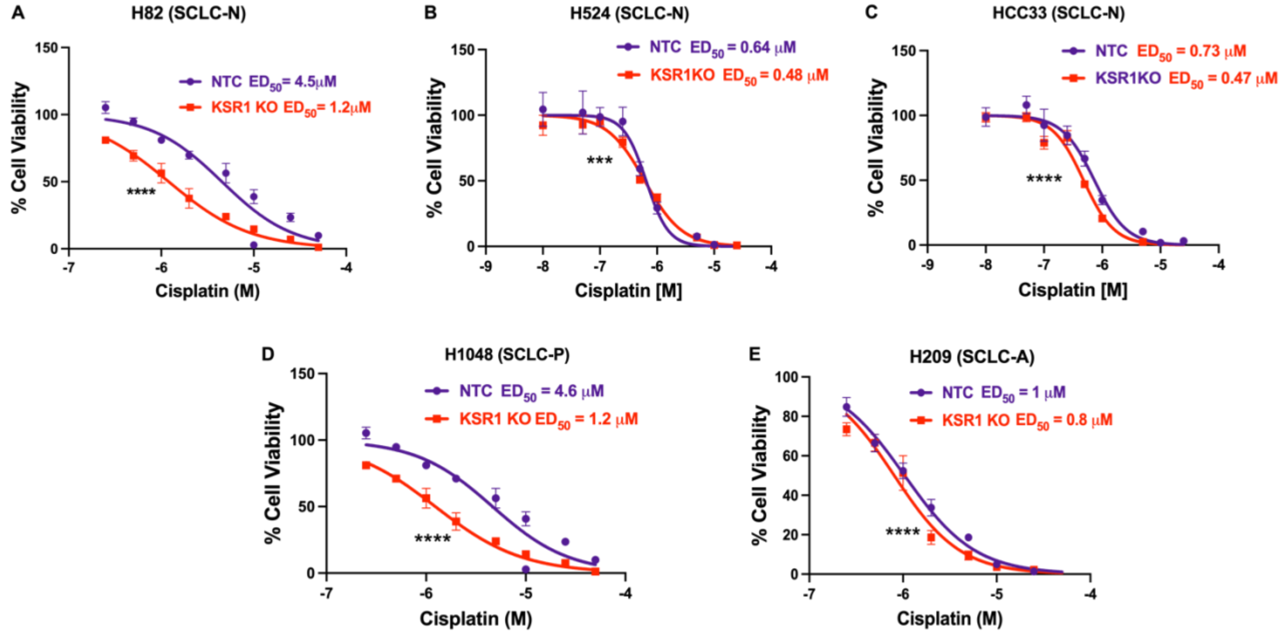


**Supplementary Fig. 3. KSR1 regulates cisplatin resistance in SCLC cell lines.** Cisplatin dose response curve with non-targeting control (NTC), and KSR1 knockout (KSR1 KO) H82 **(A)**, H524 **(B)**, HCC33 **(C)**, H1048 **(D)** and H209 **(E)** cells. ***, $p<0.001$; ****, $p<0.0001$.

**Supplementary Fig. 4.**

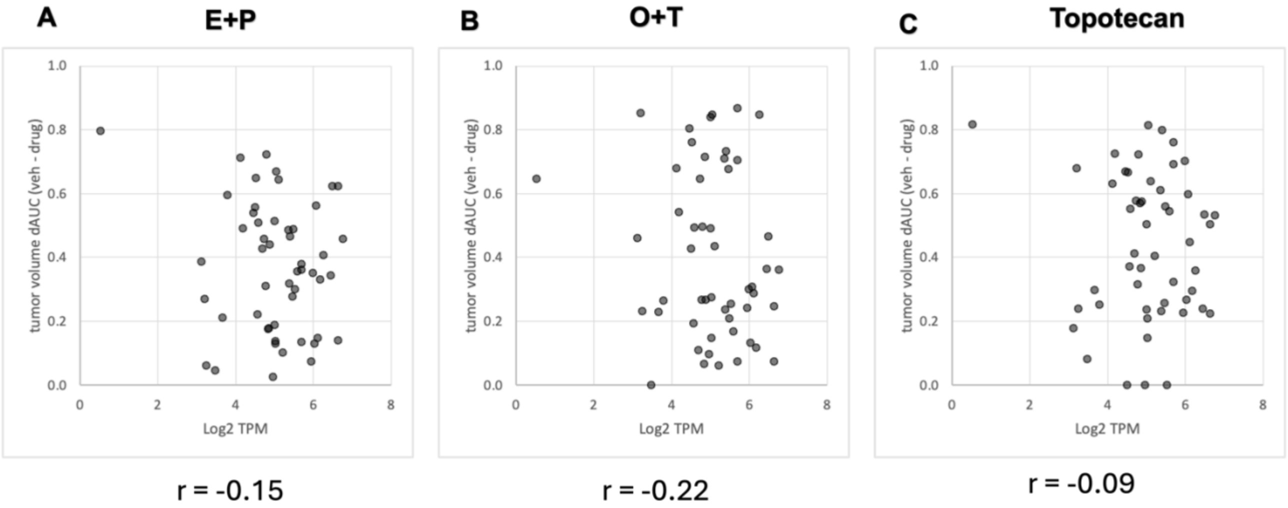


**Supplementary Fig. 4. SCLC PDXs show varying KSR1 expression in response to chemotherapy.** Correlation between average delta-AUC (drug response to chemotherapy measured from the area under tumor volume curve) and KSR1 expression in 51 PDXs under different treatment regimens as shown in **(A)** etoposide + cisplatin (E+P) **(B)** olaparib + temozolomde (O +T) **(C)** Topotecan

**Supplementary Fig. 5.**

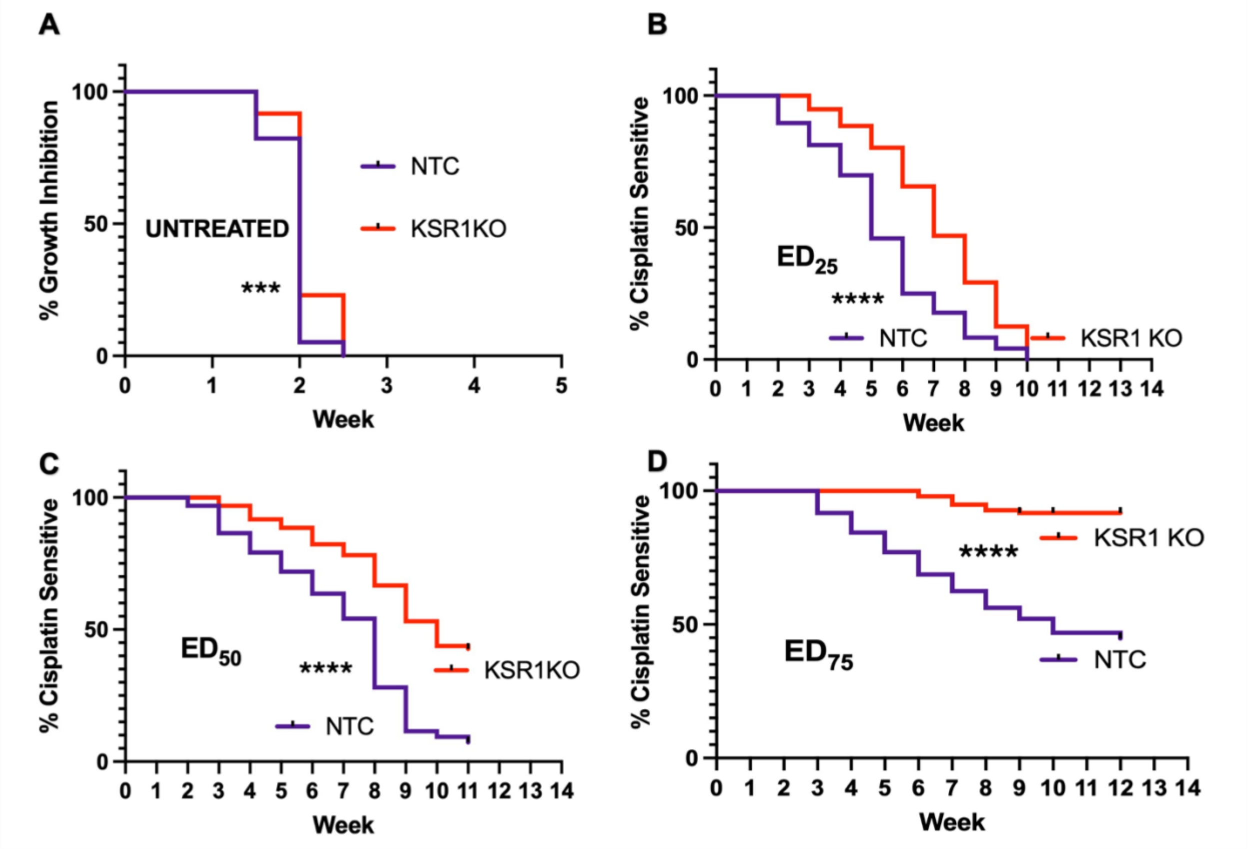


**Supplementary Fig. 5. KSR1 knockout sensitizes HCC33 cells to cisplatin in a dose-dependent manner.** Multi-well resistance assay of **(A)** untreated NTC and KSR1 KO HCC33 cells or cells treated continuously with **(B)** ED25, **(C)** ED50, or **(D)** ED75 concentrations of cisplatin. A representative experiment for each condition completed in triplicate is shown. ****, $p<0.0001$.

| Group | Cell line tested | Reference | Chiseq | P Value |
|---|---|---|---|---|
| NTC vs KSR1 KO | H82 (SCLC-N) | Fig. 2A | 14.5 | 0.000142 |
| NTC vs KSR1 KO | H524 (SCLC-N) | Fig. 2B | 24.3 | $8.45e^{-7}$ |
| NTC vs KSR1 KO | HCC33 (SCLC-N) | Fig. 2C | 23 | $1.58e^{-6}$ |
| NTC vs KSR1 KO | H526 (SCLC-P) | Fig. 2D | 14.5 | 0.000143 |
| NTC vs KSR1 KO | H1048 (SCLC-P) | Fig. 2E | 25.8 | 3.76e-7 |
| NTC vs KSR1 KO | H209 (SCLC-A) | Fig. 2F | 18.9 | 1.4e-5 |
| NTC vs KSR1 KO | H2107 (SCLC-A) | Fig. 2G | 19.3 | 1.1e-5 |
| NTC vs KSR1 KO vs KSR1AAAP vs WTKSR1 | H82 (SCLC-N) | Fig. 3C | 34.8 | 1.34e-7 |
| NTC vs ERKi | H82 (SCLC-N) | Fig. 3F | 0.117 | 0.732 |
| NTC vs ERKi | HCC33 (SCLC-N) | Fig. 3H | 0.518 | 0.472 |
| NTC vs ERKi | H1048 (SCLC-P) | Fig. 3I | 0.506 | 0.477 |
| NTC vs ERKi | H209 (SCLC-A) | Fig. 3J | 1.51 | 0.219 |
| NTC vs KSR1 KO | H82 (SCLC-N) **(in vivo)** | Fig. 4 | 7.66 | 0.00564 |
| NTC and KSR1 KO groups compared +/- Cisplatin | H82 (SCLC-N) | Fig. 5A | 302 | 3.32e-65 |
| NTC and KSR1 KO groups compared +/- Cisplatin across different doses | H524 (SCLC-N) | Fig. 5B | 552 | 2.42e-90 |
| NTC and KSR1 KO groups compared +/- Cisplatin across different doses | HCC33 (SCLC-N) | Fig. 5C | 526 | 2.57e-109 |
| NTC and KSR1 KO groups compared +/- Cisplatin across different doses | H1048 (SCLC-P) | Fig. 5D | 182 | 1.6e-37 |
| NTC and KSR1 KO groups compared +/- Cisplatin across different doses | H209 (SCLC-A) | Fig. 5E | 139 | 3.33e-28 |

**Supplementary Table 1. Statistical significance of ELDAs.** The table shows the chi squared test values (ChiSeq) and the P Values across different comparison groups for all the in vitro/ in vivo ELDAs conducted in this study.
